# Longitudinal trajectories of brain age in young individuals at familial risk of mood disorder

**DOI:** 10.1101/537951

**Authors:** Laura de Nooij, Mathew A. Harris, Emma L. Hawkins, Xueyi Shen, Toni-Kim Clarke, Stella W.Y. Chan, Tim B. Ziermans, Andrew M. McIntosh, Heather C. Whalley

## Abstract

**Background:** Within young individuals, mood disorder onset may be related to changes in trajectory of brain structure development. To date, however, longitudinal prospective studies remain scarce, showing partly contradictory findings, with a lack of emphasis on changes at the level global brain patterns. Cross-sectional adult studies have applied such methods and show that mood disorders are associated with differential aging trajectories, or accelerated brain ageing. Currently, it remains unclear therefore whether young individuals show differential brain structure ageing trajectories associated with onset of mood disorder and/or presence of familial risk.

**Methods:** Participants included young individuals (15-30 years, 53% F) from the prospective longitudinal Scottish Bipolar Family Study (SBFS) with and without a close family history of mood disorder. All were well at time of recruitment. Implementing a structural MRI-based brain age prediction model we globally assessed individual trajectories of age-related structural change with use of the difference between predicted brain age and chronological age (brain-predicted age difference; brain-PAD) at baseline and at two-year follow-up. Based on follow-up clinical assessment, individuals were categorised into three groups: (i) controls who remained well (C-well, *n* = 93), (ii) high familial risk who remained well (HR-well, *n* = 74) and (iii) high familial risk who developed a mood disorder (HR-MD, *n* = 35).

**Results:** At baseline, brain-PAD was comparable between groups. Results showed statistically significant negative trajectories of brain-PAD between baseline and follow-up for HR-MD versus C-well (β = - 0.60, *p*_*corrected*_ < .001) and HR-well (β = −0.36, *p*_*corrected*_ = .02), with a potential intermediate trajectory for HR-well (β = −0.24 years, *p*_*corrected*_ = .06).

**Conclusions:** These findings suggest that within young individuals, onset of mood disorder and familial risk may be associated with a deceleration in brain structure ageing trajectories. Extended longitudinal research will need to corroborate findings of emerging maturational lags in relation to mood disorder risk and onset.

## 1. Introduction

Mood disorders are amongst the most common psychiatric disorders, with a life-time prevalence of around 15% (Kessler and Bromet, 2013). Globally, they are the greatest contributor to non-fatal ill-health (World Health Organization, 2017). However, underlying biological mechanisms remain unclear. It is known that mood disorders are highly heritable and share complex genetic architecture; individuals with a family history of Bipolar Disorder (BD) have >10-fold increased risk of developing BD or Major Depressive Disorder (MDD) (Smoller and Finn, 2003). Mood disorders often manifest during adolescence and young adulthood (De Girolamo et al., 2012). During these life stages, age-related changes in brain structure contribute to cognitive development but also increase vulnerability to mental illness, including mood disorders (Andersen, 2003; Dahl, 2004).

From adolescence onward, decreases in brain grey matter and fine-tuning/stabilisation of synapses parallel changes in cognition and affect regulation (Giorgio et al., 2010; Spear, 2000). For higher-order cortical areas these structural trajectories extend into young adulthood (Gogtay et al., 2004; Wierenga et al., 2016, 2014). Previous prospective longitudinal studies including young individuals have shown inconsistent findings with regard to brain structure changes and mood disorder onset (Bos et al., 2018; Ducharme et al., 2014; Papmeyer et al., 2016, 2015; Whittle et al., 2014). The most consistent results suggest that the frontal cortex (Ducharme et al., 2014; Papmeyer et al., 2015) and subcortical volumes (Whittle et al., 2014) show decelerated brain structure ageing trajectories. Theoretically, decelerated trajectories may contribute to vulnerability to mood disorders, particularly when frontal-limbic brain systems of cognitive control and emotional stability are affected. Previous studies in this field mostly focused on specific regions of interest, so that spatially unbiased comprehensive approaches investigating global patterns are currently lacking. We were therefore interested in determining from longitudinal prospective data of young individuals, whether a comprehensive and spatially unconstrained measure of brain structure ageing trajectory across the brain was related to concurrent mood disorder onset and/or to familial risk.

A new framework that allows for global assessment of age-related patterns of structural change in the brain involves the estimation of an individual’s “biological brain age” from an MRI scan. Subsequent comparison with chronological age provides the brain-predicted age difference (brain-PAD) as a cross-sectional measure of brain ageing. Conceptually, when brain ageing trajectories that shape cognition and behaviour show individually different temporal dynamics, brain-PAD is expected to relate to relevant outcomes. Indeed, previous cross-sectional research within early-old aged adults showed that older appearing brains were associated with age-related diseases and mental illness (for overview see Cole et al., 2019), including mood disorders (Han et al., 2019; Koutsouleri et al., 2014; but no effect in Nenadić et al., 2017), and were furthermore predictive of mortality (Cole et al., 2018). Interestingly, accelerated brain ageing in mood disorders is in accordance with accelerated biological ageing (Rizzo et al., 2014; Sibille, 2013; Wolkowitz et al., 2011) as well as increased risk of age-related disease and mortality (e.g., Mezuk et al., 2008; Ösby et al., 2001; Pan et al., 2011).

Within longitudinal designs including younger individuals, brain-PAD has potential to show the temporal origin of accelerated brain ageing trajectories that have been observed in adult samples. Importantly, the brain-PAD approach has previously been applied and validated within samples of children and adolescents (Franke et al., 2012). Furthermore, a previous cross-sectional study did not find differences in brain-PAD between young adults at high familial risk with mood disorder diagnosis, those at high familial risk who were well, and control subjects (Hajek et al., 2017). To our knowledge, however, the current study is the first longitudinal study to apply brain-PAD methods within a sample of young individuals to investigate associations between mood disorder risk and onset, and age-related changes in brain structure.

Specifically, the current study investigated divergence of normative brain structure ageing trajectories in young individuals by applying the brain-PAD framework within a prospective longitudinal design, starting before mood disorder onset. We used data from the Scottish Bipolar Family Study (SBFS), which included young individuals who were all initially well and some of whom had a close family history of BD. Within this cohort our group previously identified differences in cortical thickness trajectories associated with high risk and mood disorder onset, including increased thickness of the left inferior frontal gyrus and left precentral gyrus in those at high risk who subsequently developed mood disorder versus cortical thickness reductions in those who remained well (Papmeyer et al. 2015a). By contrast, no subcortical volume markers of risk and illness were found (Papmeyer et al. 2016).

Investigation of white matter structure at baseline furthermore revealed reduced white matter integrity associated with familial risk (Sprooten et al. 2011), and follow-up data suggested that this finding was related to sub-clinical symptoms rather than predictive of clinical outcome (Ganzola et al. 2018). The current study builds on previous research within the BFS cohort, which identified differences in specific grey matter regions and white matter abnormalities, by investigating global trajectories of grey matter structure associated with familial risk and onset of mood disorder.

Recognising similarities between BD and MDD in symptomatology and genetic architecture, as well as the difficulty of defining a definitive stable diagnosis at young age, early-onset mood disorder was defined as having an onset of MDD or BD during adolescence or young adulthood. The longitudinal character of the study enabled the investigation of brain-PAD over two years, to assess differential brain structure ageing trajectories for those who were at high risk for mood disorder and/or subsequently developed illness.

Based on previous research, we predicted that mood disorder onset in youth would be associated with differential trajectories of brain structural change, without a specific hypothesis relating to the direction of this effect. Previous results from longitudinal developmental studies are inconclusive, but show weak evidence of decelerated trajectories, which also corresponds to theoretical developmental perspectives. Conversely, early adulthood may represent the temporal origin of the accelerated brain ageing observed in older adults, in which case we would expect an effect of mood disorder in this direction instead. We also hypothesised that the presence of familial risk would be associated with differences in brain-PAD trajectory.

## 2. Materials and Methods

### 2.1 Participants

Participants were adolescents and young adults (N = 283, age 15-30 years) recruited as part of the Scottish Bipolar Family Study (SBFS) (Chan et al., 2016; Ganzola et al., 2018; Sprooten et al., 2011; Whalley et al., 2015). Participants at high familial risk of mood disorder (HR-participants) had at least one first-degree relative or two second-degree relatives with BD type-I, and were thus at increased risk of developing a mood disorder (i.e., over 10-fold increased risk for both BD and MDD) (Smoller and Finn, 2003). Unrelated control participants without family history of BD or other mood disorder were recruited from the social networks of HR-participants, and were matched to the HR-group by age and sex. Details of familial structure within the groups are described in Appendix A1. Exclusion criteria ensured that, at the time of recruitment, all participants had no personal history of MDD, mania or hypomania, psychosis, or any other major neurological or psychiatric disorder, substance dependence, learning disability, or head injury that included loss of consciousness, and that they were without contraindications to MRI. Therefore, all individuals (HR and control) were considered well at the baseline imaging assessment.

The following additional exclusion criteria were applied in the context of the current study: (i) missing MRI or age data (*n* = 40), (ii) scans of insufficient image or segmentation quality (*n* = 15) (iii) unclear or other psychiatric diagnosis without mood disorder (*n* = 5) (see Appendix A.2), and (iv) high familial risk for mood disorder without follow-up measurement (*n* = 9). These criteria excluded 69 participants, reducing the sample size to a total of 214 participants at timepoint 1 (108 HR-participants), with follow-up timepoint 2 data available for 133 of these participants (78 HR-participants).

### 2.2 Procedure

Participants of the SBFS were invited every two years for a total of four assessments over six years (Whalley et al., 2015). Participants were interviewed and screened with the Structured Clinical Interview for DSM-IV Axis-I Disorders (SCID) (First et al., 2002) by two trained psychiatrists at timepoint 1 to ensure that they were all initially well, and at timepoint 2 to determine the presence of any mood disorder meeting diagnostic criteria since the previous assessment. Timepoint 2 clinical information was available for 93% of the included control participants, and for all included HR-participants. Of note, control participants with missing clinical information at timepoint 2 (7%) also had missing timepoint 2 MRI data, but their baseline data was retained to increase the training sample size and contributed to statistical modelling of the mean brain-PAD at baseline. Using the same procedure as previous studies on this cohort (Whalley et al. 2015; Chan et al. 2016), participants were categorised as well or diagnosed with mood disorder according to available clinical information. Individuals with well outcomes at the earlier two assessments were assumed to have remained well in the absence of further clinical information to the contrary at timepoint 3 (see Appendix A.3, Table S1). Additionally however, if individuals were subsequently found to have been diagnosed with mood disorder at further assessments (*n* = 13), they were then categorised in the mood disorder group. Including these participants in the mood disorder group enables the investigation of early disease mechanisms, while keeping the well-groups as pure as possible. Group categorisation resulted in the following groups: control participants who remained well (C-well, *n* = 93), HR-participants who remained well (HR-well, *n* = 74), and HR-participants who developed a mood disorder (HR-MD, *n* = 35, including 6 BD). As only a small number of control participants developed a mood disorder (*n* = 12, including 2 BD), these participants were not included in the main analysis. Thus, the sample for our main analysis consisted of 202 participants at baseline and 124 participants at follow-up.

The National Adult Reading Test (NART) (Nelson and Willison, 1991) and Hamilton Rating Scale for Depression (HRSD) (Hamilton, 1960) were administered at the time of scanning. The participant’s age at the time of each assessment was registered in years with a precision of two decimals. Assessments at timepoint 1, timepoint 2 and timepoint 4 included an MRI session, although only MRI measurements at timepoint 1 and timepoint 2 were considered within this study to restrict to a single scanner. The SBFS was approved by the Research Ethics Committee for Scotland, and written informed consent including consent for data linkage via medical health records was acquired from all participants.

### 2.3 MRI acquisition and pre-processing

Timepoint 1 and timepoint 2 MRI sessions were carried out on a 1.5 T Signa scanner (GE Medical, Milwaukee, USA) at the Brain Research Imaging Centre in Edinburgh and included a structural T1 weighted sequence (180 contiguous 1.2 mm coronal slices; matrix = 192 x 192; fov = 24 cm; flip angle 8°).

Pre-processing of T1 weighted scans was done in Statistical Parametric Mapping (SPM) version 12. The Computational Anatomy Toolbox (CAT) toolbox (version CAT12.3 (r1318); Gaser and Dahnke, 2018), which runs on SPM12 software, was used to segment T1-weighted MRI scans into different tissue types (for details see Appendix A.4). A cross-sectional segmentation approach was utilised in order to maximise the size of our training sample, and the longitudinal aspect of our data was handled with repeated measures linear mixed modelling (see section 2.5). This approach avoided the exclusion of participants with incomplete MRI data. CAT12 Quality Assurance metrics were used in combination with manual checks to achieve an objective and comprehensive procedure to exclude scans with artefacts or of otherwise insufficient quality (see Appendix A.4). Subsequently, modulated grey matter maps (GMM) were smoothed with a Gaussian kernel (FWHM = 8 mm). After loading the smoothed GMM (sGMM) into Python version 3.5.4, voxels were resampled into voxels of double the original voxel size, i.e. 3 x 3 x 3 mm^3.^ This reduced the number of voxels without further loss of spatial information. The sGMM were then masked with a threshold of 0.01 to ensure that voxels outside the brain were represented by value zero. The resulting sGMM were used as input for the brain-PAD model.

### 2.4 Brain-PAD model

To initially train the brain age prediction model, the training sample included all control and HR-participants that remained well (*n* = 167) in order to maximise the healthy sample size (a model including control participants only was considered underpowered, see Appendix A.5). The current model was equally balanced across timepoint 1 and timepoint 2 measurements in order to maximise the age range. Specifically, each well-group participant provided one scan for the training sample: 48 timepoint 1 scans and 46 timepoint 2 scans (i.e. all available scans) for C-well, together with 37 scans per timepoint for HR-well. HR-well timepoint 2 scans were selected based on the highest chronological ages at follow-up so that the age range covered by the training sample was maximal (*M*_*age*_ = 22.37, *SD*_*age*_ = 2.94, age range = 15.2-28.1 years; for age distributions see Figure S1 in Appendix A.6). Specifically, each well-group participant provided one scan for the training sample; this was a timepoint 2 scan for all 46 C-well participants with follow-up scan and the 37 HR-well participants with the highest chronological ages at follow-up.

Similar to previous studies (see Cole et al., 2019), the sGMM and corresponding chronological ages of the training sample were used to train a brain age prediction model. This model was implemented in Python (version 2.6.6). Corresponding to recent recommendations (Smith et al., 2019), this model initially consisted of dimension reduction of all sGMM voxels to 73 brain components (based on eigenvalue > 1) using principal component analysis (PCA) based on singular value decomposition (SVD) from scikit-learn (Pedregosa et al., 2011). We subsequently used these brain components (X) and chronological age (y) as input for estimating a Relevance Vector Regression (RVR) model with linear kernel (Tipping, 2001); this was implemented using the publicly available scikit-rvm package (https://github.com/JamesRitchie/scikit-rvm). The RVR algorithm was chosen because kernel-based methods have been most commonly implemented in brain age models (Cole et al., 2019), because linear RVR was found to be the favourable algorithm in a previous brain-PAD study (Franke et al., 2010) and because RVR does not require estimation of hyperparameters using cross-validation (a procedure that would limit our sample size).

The trained model was then applied to each participant’s sGMM to predict their brain age, ensuring that the participant for whom the brain age was being predicted was left out of the training sample to prevent bias (leave-one-out training). A residuals approach was used to regress out chronological age and gender, and subsequently calculate brain-PAD (for details see Appendix A.7), i.e. the gap between brain age prediction and chronological age. This residuals based approach is typically used to derive measures of accelerated ageing (e.g. epigenetic ageing; Chen et al., 2016; Horvath, 2013) and is recommended for the brain-PAD approach (Smith et al., 2019).

Regarding brain development, a positive brain-PAD reflected a brain-predicted age older than the chronological age of the participant, while a negative brain-PAD indicated a brain-predicted age younger than the participant’s chronological age. Changes in brain-PAD over time indicated a relative acceleration in brain maturation if brain-PAD became more positive (or less negative), or a relative deceleration in brain maturation if brain-PAD became more negative (or less positive).

Given the aim of the current study to specifically investigate brain structure ageing trajectories within the SBFS cohort, as well as the demographics of our cohort (particularly the narrow age range, also including late adolescence), we achieved within-sample model evaluation based on the brain age predictions for the training sample, using leave-one-out cross-validation.

### 2.5 Comparison of brain maturation trajectories

Since the objective of this study was to investigate deviation of brain maturation trajectories in young individuals at high risk for mood disorder and the association with illness onset, participants were divided in three groups based on clinical information as described above. Clinical information from all available assessments was considered in group categorisation as described above.

In order to compare brain structure ageing trajectories between groups, we applied a linear mixed model (LMM) to the brain-PAD measures, taking into account loss to follow-up as well as individual and family-related effects (Gueorguieva and Krystal, 2004). This was modelled using R (version 3.2.3) package nlme (Pinheiro et al. 2015) with the formula: *‘Brain-PAD ~ Timepoint * Group, random = ~1 | FamilyID / SubjectID’.* Within this single pre-defined LMM model, we were interested in the following contrasts: group differences in brain-PAD at baseline (group effect), differential trajectories of brain-PAD between groups (group by timepoint interaction effect), and group differences at follow-up. For these contrasts we tested all three pairwise comparisons, and we multiple comparison corrected results (*n* = 3 pairwise comparisons) with the Holm-Bonferroni method (Holm, 1979) using R package ‘emmeans’ (https://github.com/rvlenth/emmeans).

Exploratory analyses were conducted to further explore group differences in Brain-PAD trajectories. Firstly, we tested a longitudinal model that considered the interaction effect between age (at baseline) and group on the difference in brain-PAD between baseline and follow-up; this was modelled in R using the formula: ‘*Brain-PAD_difference ~ Age_baseline * Group, random = ~1 | FamilyID/SubjectID’*. A second exploratory analysis also modelled the brain-PAD trajectory for the group of control participants who developed a mood disorder (C-MD) within the LMM of the main analysis, considering the pairwise comparisons with control group C-well. In all of the analyses described above, continuous variables (brain-PAD, age) were transformed to Z-scores to retrieve standardised β-coefficients.

## 3. Results

### 3.1. Demographic and clinical variables

Sample sizes, demographic information and clinical measures are presented in Table 1. There were no significant differences between groups with regard to age at either timepoint, and no differences in gender, handedness and NART intelligence quotient score.

**Table 1.** Demographic and clinical characteristics.

|  | C-well | HR-well | HR-MD | p-value |
| --- | --- | --- | --- | --- |
| *n*, timepoint 1 | 93 | 74 | 35 |  |
| *n*, timepoint 2 | 46 | 47 | 31 |  |
| Age, timepoint 1<sup>a</sup> | 21.6 (3.8) | 21.6 (4.9) | 21.4 (5.2) | .82 |
| Age range | 16.3-25.6 | 15.2-26.6 | 16.0-30.0 |  |
| Age, timepoint 2<sup>a</sup> | 23.4 (3.6) | 23.8 (5.2) | 23.7 (5.0) | .45 |
| Age range | 18.3-27.6 | 17.6-28.1 | 18.1-28.1 |  |
| Gender<sup>b</sup> |  |  |  | .51 |
| Male | 42 (45.2%) | 38 (51.4%) | 14 (40.0%) |  |
| Female | 51 (54.8%) | 36 (48.6%) | 21 (60.0%) |  |
| Handedness<sup>b</sup> |  |  |  | .32 |
| Left | 5 (5.4%) | 7 (9.5%) | 2 (5.7%) |  |
| Right | 86 (92.5%) | 65 (87.8%) | 33 (94.3%) |  |
| Mixed | 0 (0%) | 2 (2.7%) | 0 (0%) |  |
| Unknown | 2 (2.2%) | 0 (0%) | 0 (0%) |  |
| NART score<sup>a</sup> | 111 (9.0) | 111 (10.7) | 107 (8.9) | .15 |
| HRSD, timepoint 1<sup>a</sup> | 0 (1.0) | 0 (2.0) | 2 (5.5) | <.001***<sup>a</sup> |
| HRSD, timepoint 2<sup>a</sup> | 0 (2.0) | 0 (1.0) | 5 (8.0) | <.001***<sup>a</sup> |

\*\*\* $p < .001$
Note: Individual NART scores were averaged over all completed assessments (max. 4).
<sup>a</sup> Medians and interquartile ranges for variables not normally distributed (Kruskal-Wallis test)
<sup>b</sup> Frequency and percentages for categorical variables (Chi-squared test).
<sup>a</sup> HR-MD vs. C-well and HR-MD vs. HR-well (Chi-squared test, followed by Dunn's test for pairwise comparisons)
C-well, group of participants without family history who remained well; HR-MD, group of participants at high familial risk who developed a mood disorder; HRSD, Hamilton Rating Scale for Depression; HR-well, group of participants at high familial risk who remained well; NART, National Adult Reading Test.

However, HR-MD participants reported greater depression symptomatology on the HRSD as compared to the groups of participants who remained well (C-well and HR-well) at both timepoints (Table 1). At baseline, seven HR-MD participants (20%; *M*_HRSD_ = 11.4) reported subclinical symptoms of depression (defined as HRSD score > 7). At timepoint 2, ten HR-MD participants reported symptoms of depression (defined as HRSD score > 7). For two of these participants depression symptoms were at subclinical level, as they were not yet diagnosed with a mood disorder. In contrast, there was a very low prevalence of subclinical depression symptomatology within the well-groups: two participants at timepoint 1 (1.2%; *M*_HRSD_ = 9.0) and two other participants (with included follow-up scans) at timepoint 2 (2.6%; *M*_HRSD_ = 9.0).

Participants did not report any use of psychotropic medication at baseline. At follow-up, six included HR-MD participants reported the use of psychotropic medication, whereas none of the well-group participants received medication for the treatment of psychiatric symptoms.

HR-MD showed lower attrition (11.4%) than C-well (50.5%; χ2(1) = 14.6, *p* < .001) and HR-well (36.5%; χ2(1) = 6.2, *p* = .01), with no significant difference between C-well and HR-well (χ2(1) = 2.8, *p* = .10). None of the clinical or demographic variables at baseline differed between those individuals with and without a follow-up scan (see Table S2 in Appendix B.1).

### 3.2 Model evaluation

Our model showed a significant positive Pearson correlation between predicted brain age and chronological age (*r*(165) = .40, *p* < .001), and a mean absolute error (MAE) of 2.21 years (scaled MAE = MAE / age range = 0.17; see Appendix B.2) within the training sample. For a discussion on model evaluation within the context of the current study, see section 4 (discussion) and Appendix B.2.

The 73 brain components that were used as input for the brain age prediction algorithm indicated a mean total explained variance of 84.0% (*SD* = 0.0004) for all (leave-one-out) training sample dimension reduction iterations. These brain components showed loadings distributed across the brain, because dimension reduction was spatially unconstrained. This complicated unbiased interpretation (Smith et al., 2019), and therefore, also given our aim to comprehensively assess global patterns of brain structure ageing trajectories, these components were not further explored. However, we do present visualisation of these brain components in order to illustrate the method (Figure S3 in Appendix B.3).

### 3.3 Comparison of brain maturation trajectories

#### 3.3.1 Comparison at baseline

Group allocation based on diagnostic information resulted in mean brain-PADs of +0.04 (*SD* = 1.14, *n* = 93) for C-well, −0.36 (*SD* = 1.22, *n* = 74) for HR-well, and −0.01 (*SD* = 1.39, *n* = 35) for HR-MD. Results of the LMM (Table 2) suggested lower baseline brain-PAD for HR-well compared to C-well (−0.42 years; β = −0.37, *p* = .03, *p*_*corrected*_ = .08), but statistical significance did not survive multiple comparison correction. There were no baseline differences in brain-PADs for HR-MD versus C-well (−0.05 years; β = −0.07, *p*_*corrected*_ = .73) or HR-MD versus HR-well (0.35 years; β = 0.30, *p*_*corrected*_ = .24).

**Table 2.** Fixed effects of linear mixed model applied to investigate group differences in the brain-predicted age difference (brain-PAD).

| Main model - Comparison to C-well |  |  |  |  |  |
| --- | --- | --- | --- | --- | --- |
| Fixed effect | β-coefficient | SE | df | t-value | p-value |
| (Intercept) | 0.09 | 0.10 | 174 | 0.84 | .55 |
| Timepoint 2 | 0.37 | 0.09 | 121 | 4.10 | <.001*** |
| HR-well | -0.37 | 0.16 | 25 | -2.33 | .03* |
| HR-MD | -0.07 | 0.20 | 25 | -0.35 | .72 |
| Timepoint 2*HR-well | -0.24 | 0.13 | 121 | -1.93 | .06 |
| Timepoint 2*HR-MD | -0.60 | 0.14 | 121 | -4.23 | <.001*** |
| Main model - Comparison to HR-well |  |  |  |  |  |
| Fixed effect | β-coefficient | SE | Df | t-value | p-value |
| HR-MD | 0.30 | 0.21 | 25 | 1.14 | .16 |
| Timepoint 2*HR-MD | -0.36 | 0.14 | 121 | -2.52 | .01* |

\* $p < .05$ , \*\* $p < .01$ , \*\*\* $p < .001$
Note: β-coefficients are standardised following scaling of the outcome variable.
C-well, group of participants without family history who remained well; HR-MD, group of participants at high familial risk who developed a mood disorder; HR-well, group of participants at high familial risk who remained well.

#### 3.3.2 Brain structure ageing trajectories

Results showed a statistically significant timepoint by group interaction effect for HR-MD compared to C-well (−0.70 years; β = −0.60, *p*_*corrected*_ < .001) and HR-well (−0.43 years; β = −0.36, *p*_*corrected*_ = .02), indicating decelerating brain structure ageing trajectories. Besides that, HR-well showed a intermediate trajectory (−0.28 years; β = −0.24, *p*_*corrected*_ = .06) which was not statistically significant. Figure 1 displays brain maturation trajectories per group as modelled by unstandardised LMM fixed effects; for clarity, these trajectories are displayed *relative to the control group* following correction for the effects observed in C-well (i.e., intercept and the significant timepoint coefficient, see Table 2). Figure 2 shows the heterogeneity in observed brain maturation trajectories by displaying the participants’ individual changes in brain-PAD over time.

**Figure 1.**
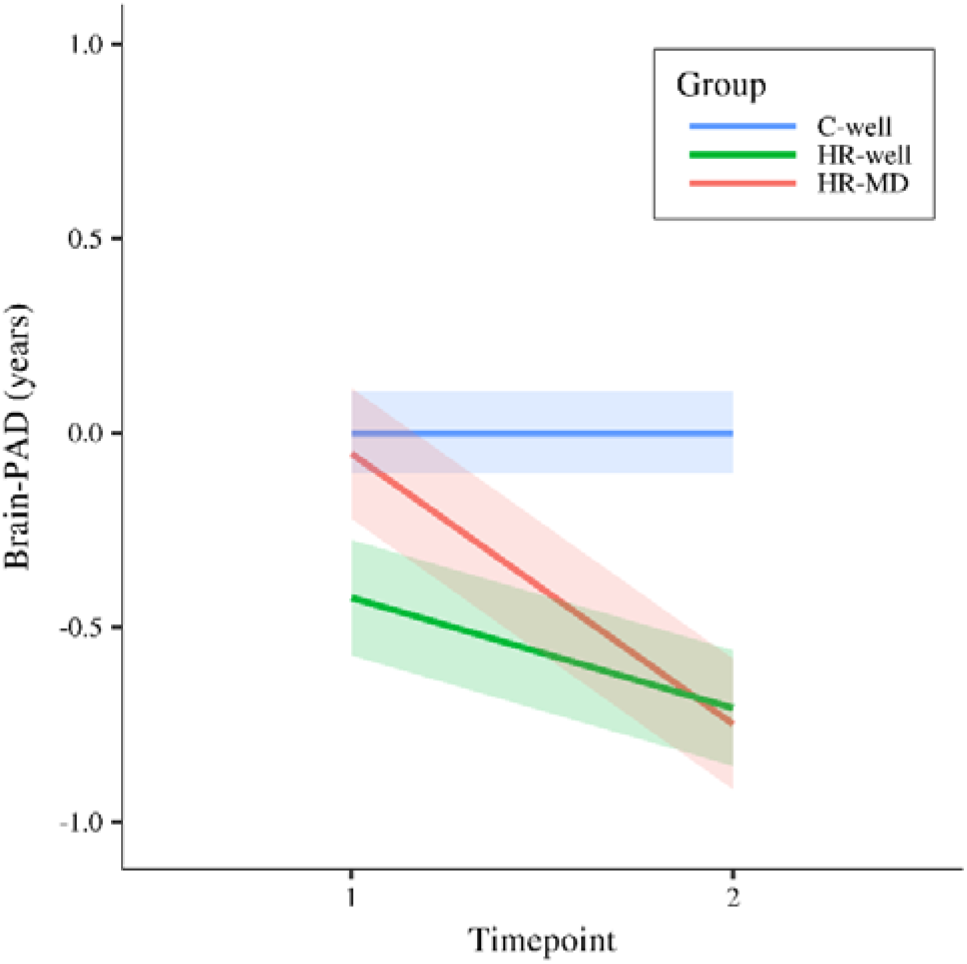
Modelled fixed effects of the brain-predicted age difference (brain-PAD) per group, for clarity corrected for effects in C-well (i.e., the intercept and timepoint coefficients) as this group functions as control group. Shaded areas display standard errors of the timepoint by group interaction effects. Brain-PAD, brain-predicted age difference; C-well, group of participants without family history who remained well; HR-MD, group of participants at high familial risk who developed a mood disorder; HR-well, group of participants at high familial risk who remained well.

**Figure 2.**
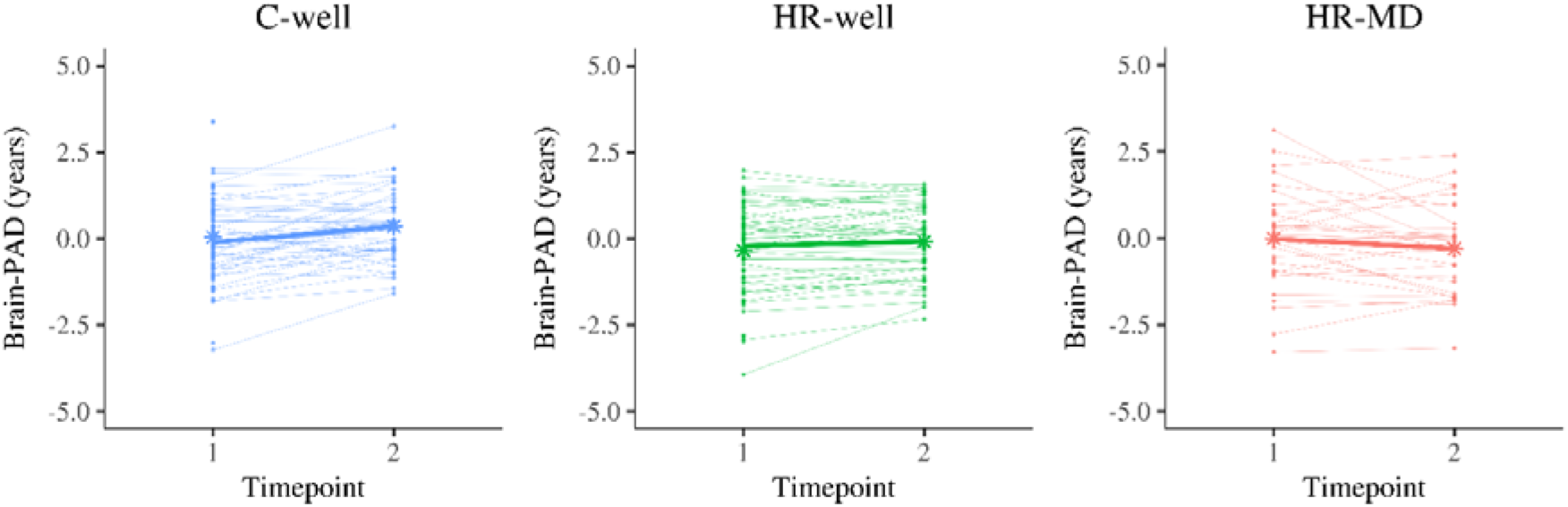
Display of brain structure ageing trajectories per participant, reflecting a changing brain-predicted age difference (brain-PAD) between timepoint 1 and timepoint 2 (two years apart). Each panel contains the trajectories of one group in thin line graphs, whereas the thicker line graph represents the average trajectory of that group (of complete cases). The star dots display the mean brain-PAD at each timepoint. Left panel, C-well; Middle panel, HR-well; Right panel, HR-MD. Brain-PAD, brain-predicted age difference; C-well, group of participants without family history who remained well; HR-MD, group of participants at high familial risk who developed a mood disorder; HR-well, group of participants at high familial risk who remained well.

#### 3.3.3 Comparison at follow-up

At follow-up, two years later, the mean brain-PADs were +0.36 (*SD* = 1.01) for C-well, −0.08 (*SD* = 1.04) for HR-well, and −0.30 (*SD* = 1.30) for HR-MD. Results indicated a statistical significant difference in brain-PAD between HR-MD and C-well (−0.69 years; β = −0.61, *p* = .02, *p*_*corrected*_ = .06), and between HR-well and C-well (−0.54 years; β = −0.48, *p* = .04, *p*_*corrected*_ = .06), although these results not survive multiple comparison correction. We found no evidence for a difference between HR-MD and HR-well at follow-up (−0.15 years; β = −0.13, *p*_*corrected*_ = .57).

#### 3.3.4 Exploratory findings

Our exploratory longitudinal model showed similar group trajectories as our main model, and furthermore suggested that within our sample, younger individuals from the HR-MD group showed greater deceleration in their structural brain trajectory than older HR-MD individuals (see Figure S4 and Table S4 in Appendix B.4). Exploratory findings of the model including the C-MD group showed a non-significant negative trajectory of brain-PAD for this group (−0.42 years; β = −0.36, *p* = .11; see Table S5 and Figure S5 in Appendix B.5).

## 4 Discussion

The results of the current study showed that in young individuals at familial risk the onset of mood disorder was associated with differences in brain structure changes over time. Statistically significant reductions in brain-PAD indicated decelerated brain structure ageing trajectories in young HR individuals who developed a mood disorder as compared to control and HR individuals who remained well. Intermediate effect sizes indicated that young individuals who were at risk but remained well showed intermediate trajectories. These results suggest genetic predisposition to mood disorder is accompanied by changes in adolescent brain structural development trajectories that are increased with the onset of mood disorder. Further research will be necessary to disentangle the role of genetic predisposition and additional environmental risk factors (e.g. adverse life events) on global age-related brain structure changes. As development of mood disorder was associated with a more decelerating trajectory, differences observed for the mood disorder group may also partly reflect prodromal symptoms or early-disease mechanisms of psychological stress. Notably, all groups showed considerable heterogeneity in the direction, size and emergence of the individual brain-PADs. Additional research is therefore required to substantiate the hypothesis that the emergence of a lag in brain structure ageing in youth indicates mood disorder onset and familial risk.

The current findings correspond with previous neuroimaging studies using different methods that also indicated deceleration, in the same as well as independent prospective longitudinal cohorts (Ducharme et al. 2014; Whittle et al. 2014; Papmeyer et al. 2015a). According to empirical-based neural models, dysfunctions in medial prefrontal networks and limbic areas underlie disturbances in emotion regulation and cognitive control (e.g. Drevets et al. 2008) which are proposed to play a causal role in the development of mood disorder (Nolen-Hoeksema et al., 2008; Phillips et al., 2008). Correspondingly, previous findings within the same cohort have revealed that illness risk and onset were associated with differential cortical thickness trajectories in prefrontal areas (Papmeyer et al. 2015a) as well as differential patterns of brain activation during emotional tasks in cortico-thalamic-limbic regions (Whalley et al. 2015; Chan et al. 2016), and neurocognitive performance was found to be a trait-marker of familial risk (Papmeyer et al. 2015b). Although the current study adopted a global approach (thus refraining from investigation of regional brain structural or functional development), we speculate about a potential neural mechanism by which decelerated trajectories of brain structural change in young individuals potentially disrupt frontal and limbic brain networks that underly emotion regulation and cognitive control, consequently increasing vulnerability to mood disorder. Inferences of causality however should be drawn with caution. It is important to consider that though prospective longitudinal studies are one approach to examining causal processes, interpretation is complex. In the current study for example, individuals who subsequently developed a mood disorder also showed higher mean subclinical depression symptomatology at baseline. This could be interpreted as a predictor of subsequent illness, or indeed as prodromal or early stages of the illness itself.

Importantly, our findings suggest disease-related brain ageing deceleration may emerge in young individuals, in contrast to findings of accelerated ageing in older adults with mood disorder (Koutsouleris et al., 2014; Sibille, 2013; Wolkowitz et al., 2011). This reflects non-linear brain ageing trajectories, which are well-established (Giedd et al., 1999; Scahill et al., 2003; Shaw et al., 2008; Tamnes et al., 2010; Wierenga et al., 2014). During adolescence and early adulthood, brain ageing represents continued development, and decelerated brain structure changes may therefore be disadvantageous In later life, brain ageing reflects degeneration, and accelerated ageing therefore also corresponds to brain structure deficits. Therefore, accelerated ageing observed in older age could also be a vulnerability factor to mood disorder disease processes (but perhaps also a negative consequence). Although our findings in young individuals are in the opposite direction to previous findings in older individuals, both indicate a poorer trajectory in brain structure changes; further research will need to determine the turning point as well as the specificity to mood disorder psychopathology.

The current study applied a novel pattern recognition method that, to our knowledge, has not been previously applied to a longitudinal cohort of young individuals at risk of mood disorder. This approach derives a global measure of brain structure, which captures the complexity of spatial and temporal dynamics of brain ageing. Challenges in collecting clinical data from young individuals mean that large cohorts are scarce; the SBFS provided a unique opportunity to investigate the dynamics of brain structure ageing trajectories in relation to mood disorder. Development of mood disorder was found to be associated with decelerated age-related changes in brain grey matter, which could not have been identified within a cross-sectional design. Clinical information was also available for up to six years, which produced some heterogeneity in the HR-MD group, due to a range in times before onset, but also provided confidence that those classified as HR-well were not in the early stages of mood disorder at the time of imaging assessments. Overall, the current study of the SBFS shows unique strengths for youth mental health research. Ongoing work on the sample seeks to implement data linkage at 10+ years to obtain more definitive, stable diagnoses.

One limitation of our study was the low correlation between brain age prediction and chronological age, reflecting suboptimal performance of the brain age prediction model. However, this is probably also related to the tight age range of our sample as well as individual differences within this life stage. Performance of the model was constraint by the limited cohort size, as for the purpose of the current study, we adopted a within-sample approach. In order to maximise the use of available data in building our brain age prediction model, we adhered to a cross-sectional segmentation approach and included all individuals who remained well, also those at high familial risk for mood disorder. Brain-PAD slightly increased over time within the control group, indicating that a brain-PAD of zero did not indicate normative brain maturation in the current study (for details see Appendix B.2). This suggests that the brain age prediction model was biased by familial risk related structural differences following inclusion of HR-well participants within the training sample. Although this explanation would not invalidate our results, as it would suggest valid comparison of relative group differences in brain-PAD, it is considered a limitation that we cannot reliably tease apart the familial risk effect from the normative trajectory. Importantly, we directly addressed potential threats to the validity of the brain age prediction model, to the extent possible within the current sample, with a deliberate pre-defined approach consisting of dimension reduction, sparse RVR modelling and a residuals approach, in order to prevent overfitting and thereby optimise the overall model validity. However, our prediction model was not validated for generalisability to other samples because of challenges related to scanner heterogeneity, so that transferability of the model remains uncertain. Further limitations of the current study are that we were unable to investigate differences between MDD and BD, and that we cannot exclude the possibility of medication effects, although use of psychotropic medications was limited within the current sample (see section 3.1). Additionally, out-of-sample predictions often show more prediction error than within-sample predictions achieved using cross-validation (Varoquaux et al. 2017). Although participants from all three groups belonged to the same cohort and were recruited and assessed according to the same procedures, brain age predictions for individuals with mood disorder onset (i.e., HR-MD) were out-of-training-sample predictions, whereas predictions for individuals who remained well (i.e., HR-well and C-well) required leave-one-out cross-validation. Although these differential prediction procedures may have led to increased prediction error for HR-MD, our finding of intermediate trajectories in the HR-well group strongly suggests that results in HR-MD are unlikely to be driven entirely by increased random error. Further, additional testing of out of training sample scans of well-group participants indicated that prediction error was not significantly increased compared to within training sample estimates (Appendix B.2), conferring further confidence in our findings. However, taken together, the findings of the current study should be interpreted in the context of these limitations.

In order to resolve the above limitations, future research should aim to replicate our results within a larger sample (Button et al., 2013; Jollans et al., 2019). Large-scaled and extended MRI follow-up assessments would furthermore allow the application of a longitudinal brain age prediction model, which will provide a more nuanced understanding of individual developmental trajectories. A sufficient sample size would also allow for investigation of MDD and BD separately, and could account for potential medication effects. Spatial interpretability of the current model’s brain age prediction was limited, but with a larger sample methods such as orthonormal projective non-negative matrix factorisation (OPNMF) could provide information about specific regions or networks involved in associations between brain age and mood disorder (Sotiras et al., 2017, 2015; Varikuti et al., 2018). Additionally, our exploratory findings suggest it may be useful to investigate associations between brain structure ageing trajectories and mood disorder in a slightly younger sample. For now, the present study lays a theoretical and empirical foundation for the field to build upon, and will hopefully encourage further longitudinal studies of clinical youth cohorts. In the future, replication and further investigation of the association between mood disorder and decelerated brain structure ageing trajectories may provide important insights into the prediction of mood disorder onset in young individuals.

## Supporting information

Supplementary Methods (Appendix A) and Supplementary Results (Appendix B) can be found in the online version of this article.

## Acknowledgements

We would like to thank all of the participants who took part in the study and the radiographers who acquired the MRI scans. We also acknowledge the support of National Health Service (NHS) Research Scotland, through the Scottish Mental Health Research Network (www.smhrn.org.uk), who provided assistance with subject recruitment and cognitive assessments, and health professionals who cooperated in accessing clinical health record information. This study was conducted at the Brain Research Imaging Centre (http://www.bric.ed.ac.uk), which is supported by SINAPSE (Scottish Imaging Network, a Platform for Scientific Excellence, http://www.sinapse.ac.uk). All imaging aspects received financial support from the Dr Mortimer and Theresa Sackler Foundation. This work was also supported by IMAGEMEND, which received funding from the European Community’s Seventh Framework Programme (FP7/2007-2013) [grant number 602450]; and Wellcome Trust Strategic Award [grant number 104036/Z/14/Z].

## Ethical standards

The authors assert that all procedures contributing to this work comply with the ethical standards of the relevant national and institutional committees on human experimentation and with the Helsinki Declaration of 1975, as revised in 2008.

## References

Andersen, S.L., 2003. Trajectories of brain development: Point of vulnerability or window of opportunity? Neurosci. Biobehav. Rev. 27, 3–18. https://doi.org/10.1016/S0149-7634(03)00005-8

Bos, M.G.N., Peters, S., van de Kamp, F.C., Crone, E.A., Tamnes, C.K., 2018. Emerging depression in adolescence coincides with accelerated frontal cortical thinning. J. Child Psychol. Psychiatry Allied Discip. 59, 994–1002. https://doi.org/10.1111/jcpp.12895

Button, K.S., Ioannidis, J.P.A., Mokrysz, C., Nosek, B.A., Flint, J., Robinson, E.S.J., Munafò, M.R., 2013. Power failure: Why small sample size undermines the reliability of neuroscience. Nat. Rev. Neurosci. 14, 365–376. https://doi.org/10.1038/nrn3475

Chan, S.W.Y., Sussmann, J.E., Romaniuk, L., Stewart, T., Lawrie, S.M., Hall, J., McIntosh, A.M., Whalley, H.C., 2016. Deactivation in anterior cingulate cortex during facial processing in young individuals with high familial risk and early development of depression: fMRI findings from the Scottish Bipolar Family Study. J. Child Psychol. Psychiatry 57, 1277–1286. https://doi.org/10.1111/jcpp.12591

Chen, B.H., Marioni, R.E., Colicino, E., Peters, M.J., Ward-Caviness, C. K., Tsai, P.C., Roetker, N.S., Just, A.C., Demerath, E.W., Guan, W., Bressler, J., Fornage, M., Studenski, S., Vandiver, A.R., Moore, A.Z., Tanaka, T., Kiel, D.P., Liang, L., Vokonas, P., Schwartz, J., Lunetta, K.L., Murabito, J.M., Bandinelli, S., Hernandez, D.G., Melzer, D., Nalls, M., Pilling, L.C., Price, T.R., Singleton, A.B., Gieger, C., Holle, R., Kretschmer, A., Kronenberg, F., Kunze, S., Linseisen, J., Meisinger, C., Rathmann, W., Waldenberger, M., Visscher, P.M., Shah, S., Wray, N.R., McRae, A.F., Franco, O.H., Hofman, A., Uitterlinden, A.G., Absher, D., Assimes, T., Levine, M.E., Lu, A.T., Tsao, P.S., Hou, L., Manson, J.A.E., Carty, C.L., LaCroix, A.Z., Reiner, A.P., Spector, T.D., Feinberg, A.P., Levy, D., Baccarelli, A., Meurs, J. van, Bell, J.T., Peters, A., Deary, I.J., Pankow, J.S., Ferrucci, L., Horvath, S., 2016. DNA methylation-based measures of biological age: Meta-analysis predicting time to death. Aging 8, 1844–1865. https://doi.org/10.18632/aging.101020

Cole, J.H., Marioni, R.E., Harris, S.E., Deary, I.J., 2019. Brain age and other bodily ‘ages’: implications for neuropsychiatry. Mol. Psychiatry 24, 266–281. https://doi.org/10.1038/s41380-018-0098-1

Cole, J.H., Ritchie, S.J., Bastin, M.E., Valdés Hernández, M.C., Muñoz Maniega, S., Royle, N., Corley, J., Pattie, A., Harris, S.E., Zhang, Q., Wray, N.R., Redmond, P., Marioni, R.E., Starr, J.M., Cox, S.R., Wardlaw, J.M., Sharp, D.J., Deary, I.J., 2018. Brain age predicts mortality. Mol. Psychiatry 23, 1385–1392. https://doi.org/10.1038/mp.2017.62

Dahl, R.E., 2004. Adolescent brain development: A period of vulnerabilities and opportunities. Ann. N. Y. Acad. Sci. 1021, 1–22. https://doi.org/10.1196/annuals.1308.001

De Girolamo, G., Dagani, J., Purcell, R., Cocchi, A., McGorry, P.D., 2012. Age of onset of mental disorders and use of mental health services: Needs, opportunities and obstacles. Epidemiol. Psychiatr. Sci. 21, 47–57. https://doi.org/10.1017/S2045796011000746

Ducharme, S., Albaugh, M.D., Hudziak, J.J., Botteron, K.N., Nguyen, T., Truong, C., Evans, A.C., Karama, S., 2014. Anxious/depressed symptoms are linked to right ventromedial prefrontal cortical thickness maturation in healthy children and young adults. Cereb. Cortex 2941–2950. https://doi.org/10.1093/cercor/bht151

First, M.B., Spitzer, R.L., Gibbon, M., Williams, J.B., 2002. Structured Clinical Interview for DSM-IV-TR Axis I Disorders, Research Version, Patient Edition with Psychotic Screen. Biometrics Research, New York State Psychiatric Institute, New York.

Franke, K., Luders, E., May, A., Wilke, M., Gaser, C., 2012. Brain maturation: Predicting individual BrainAGE in children and adolescents using structural MRI. Neuroimage 63, 1305–1312. https://doi.org/10.1016/j.neuroimage.2012.08.001

Franke, K., Ziegler, G., Klöppel, S., Gaser, C., 2010. Estimating the age of healthy subjects from T1-weighted MRI scans using kernel methods: Exploring the influence of various parameters. Neuroimage 50, 883–892. https://doi.org/10.1016/j.neuroimage.2010.01.005

Ganzola, R., McIntosh, A.M., Nickson, T., Sprooten, E., Bastin, M.E., Giles, S., Macdonald, A., Sussmann, J., Duchesne, S., Whalley, H.C., 2018. Diffusion tensor imaging correlates of early markers of depression in youth at high-familial risk for bipolar disorder. J. Child Psychol. Psychiatry 59, 917–927. https://doi.org/10.1111/jcpp.12879

Gaser, C., Dahnke, R., 2018. CAT – A Computational Anatomy Toolbox for SPM. Giedd, J.N., Blumenthal, J., Jeffries, N.O., Castellanos, F.X., Liu, H., Zijdenbos, A., Paus, T., Evans, A.C., Rapoport, J.L., 1999. Brain development during childhood and adolescence: A longitudinal MRI study. Nat. Neurosci. 2, 861–863. https://doi.org/10.1038/13158

Giorgio, A., Watkins, K.E., Chadwick, M., James, S., Winmill, L., Douaud, G., De Stefano, N., Matthews, P.M., Smith, S.M., Johansen-Berg, H., James, A.C., 2010. Longitudinal changes in grey and white matter during adolescence. Neuroimage 49, 94–103. https://doi.org/10.1016/j.neuroimage.2009.08.003

Gogtay, N., Giedd, J.N., Lusk, L., Hayashi, K.M., Greenstein, D., Vaituzis, A.C., Nugent, T.F., Herman, D.H., Clasen, L.S., Toga, A.W., Rapoport, J.L., Thompson, P.M., 2004. Dynamic mapping of human cortical development during childhood through early adulthood. Proc. Natl. Acad. Sci. 101, 8174–8179. https://doi.org/10.1073/pnas.0402680101

Gueorguieva, R., Krystal, J.H., 2004. Move over ANOVA: Progress in analyzing repeated-measures data and its reflection in papers published in the Archives of General Psychiatry. Arch. Gen. Psychiatry 61, 310–317.

Hajek, T., Franke, K., Kolenic, M., Capkova, J., Matejka, M., Propper, L., Uher, R., Stopkova, P., Novak, T., Paus, T., Kopecek, M., Spaniel, F., Alda, M., 2017. Brain Age in early stages of Bipolar Disorders or Schizophrenia. Schizophr. Bull. 190–198. https://doi.org/10.1093/schbul/sbx172

Hamilton, M., 1960. A rating scale for Depression. J. Neurol. Neurosurg. Psychiatry 23, 56–62.

Han, L.K., Dinga, R., Hahn, T., Ching, C., Eyler, L., Aftanas, L., Aghajani, M., Aleman, A., Baune, B., Berger, K., Brak, I., Filho, G.B., Carballedo, A., Connolly, C., Couvy-Duchesne, B., Cullen, K., Dannlowski, U., Davey, C., Dima, D., Duran, F., Enneking, V., Filimonova, E., Frenzel, S., Frodl, T., Fu, C., Godlewska, B., Gotlib, I., Grabe, H., Groenewold, N., Grotegerd, D., Gruber, O., Hall, G., Harrison, B., Hatton, S., Hermesdorf, M., Hickie, I., Ho, T., Hosten, N., Jansen, A., Kahler, C., Kircher, T., Klimes-Dougan, B., Kramer, B., Krug, A., Lagopoulos, J., Leenings, R., MacMaster, F., MacQueen, G., McIntosh, A., McLellan, Q., McMahon, K., Medland, S., Mueller, B., Mwangi, B., Osipov, E., Portella, M., Pozzi, E., Reneman, L., Repple, J., Rosa, P., Sacchet, M., Saemann, P., Schnell, K., Schrantee, A., Simulionyte, E., Soares, J., Sommer, J., Stein, D., Steinstrater, O., Strike, L., Thomopoulos, S., Tol, M.-J. van, Veer, I., Vermeiren, R., Walter, H., Wee, N. van der, Werff, S. van der, Whalley, H., Winter, N., Wittfeld, K., Wright, M., Wu, M.-J., Volzke, H., Yang, T., Zannias, V., Zubicaray, G. de, Zunta-Soares, G., Abe, C., Alda, M., Andreassen, O., Boen, E., Bonnin, C., Canales-Rodriguez, E., Cannon, D., Caseras, X., Chaim-Avancini, T., Elvsashagen, T., Favre, P., Foley, S., Fullerton, J., Goikolea, J., Haarman, B., Hajek, T., Henry, C., Houenou, J., Howells, F., Ingvar, M., Kuplicki, R., Lafer, B., Landen, M., Machado-Vieira, R., Malt, U., McDonald, C., Mitchell, P., Nabulsi, L., Otaduy, M., Overs, B., Polosan, M., Pomarol-Clotet, E., Radua, J., Rive, M., Roberts, G., Ruhe, H., Salvador, R., Sarro, S., Satterthwaite, T., Savitz, J., Schene, A., Schofield, P., Serpa, M., Sim, K., Soeiro-de-Souza, M., Sutherland, A., Temmingh, H., Timmons, G., Uhlmann, A., Vieta, E., Wolf, D., Zanetti, M., Jahanshad, N., Thompson, P., Veltman, D., Penninx, B., Marquand, A., Cole, J., Schmaal, L., 2019. Brain Aging in Major Depressive Disorder: Results from the ENIGMA Major Depressive Disorder working group. bioRxiv [pre-print] 1–33. https://doi.org/10.1101/560623

Holm, S., 1979. A simple sequentially rejective multiple test procedure. Scand. J. Stat. 6, 65–70.

Horvath, S., 2013. DNA methylation age of human tissues and cell types. Genome Biol. R115. https://doi.org/10.1145/2820783.2820789

Jollans, L., Boyle, R., Artiges, E., Banaschewski, T., Desrivi, S., Smolka, M.N., Walter, H., Grigis, A., Martinot, J., Schumann, G., Garavan, H., Whelan, R., 2019. Quantifying performance of machine learning methods for neuroimaging data. Neuroimage 199, 351–365. https://doi.org/10.1016/j.neuroimage.2019.05.082

Kessler, R.C., Bromet, E.J., 2013. The Epidemiology of Depression Across Cultures. Annu. Rev. Public Health 24, 119–138. https://doi.org/10.1146/annurev-publhealth-031912-114409

Koutsouleris, N., Davatzikos, C., Borgwardt, S., Gaser, C., Bottlender, R., Frodl, T., Falkai, P., Riecher-Rössler, A., Möller, H.J., Reiser, M., Pantelis, C., Meisenzahl, E., 2014. Accelerated brain aging in schizophrenia and beyond: A neuroanatomical marker of psychiatric disorders. Schizophr. Bull. 40, 1140–1153. https://doi.org/10.1093/schbul/sbt142

Mezuk, B., Eaton, W.W., Albrecht, S., Golden, S.H., 2008. Depression and type 2 diabetes over the lifespan: A meta-analysis. Diabetes Care 31, 2383–2390. https://doi.org/10.2337/dc08-0985

Nelson, H.E., Willison, J.R.T., 1991. National Adult ReadingTest (NART): Test manual, 2nd ed. Windsor: NFER-Nelson.

Nenadić, I., Dietzek, M., Langbein, K., Sauer, H., Gaser, C., 2017. BrainAGE score indicates accelerated brain aging in schizophrenia, but not bipolar disorder. Psychiatry Res. Neuroimaging 266, 86–89. https://doi.org/10.1016/j.pscychresns.2017.05.006

Nolen-Hoeksema, S., Wisco, B.E., Lyubomirsky, S., 2008. Rethinking rumination. Perspect. Psychol. Sci. 3, 400–424. https://doi.org/10.1111/j.1745-6924.2008.00088x.

Ösby, U., Brandt, L., Correia, N., Ekbom, A., Sparén, P., 2001. Excess mortality in bipolar and unipolar disorder in Sweden. Arch. Gen. Psychiatry 58, 844–850. https://doi.org/10.1001/archpsyc.58.9.844

Pan, A., Sun, Q., Okereke, O.I., Rexrode, K.M., Hu, F.B., 2011. Depression and risk of stroke morbidity and mortality: A meta-analysis and systematic review. J. Am. Med. Assoc. 306, 1241–1249. https://doi.org/10.1001/jama.2011.1282

Papmeyer, M., Giles, S., Sussmann, J.E., Kielty, S., Stewart, T., Lawrie, S.M., Whalley, H.C., McIntosh, A.M., 2015. Cortical thickness in individuals at high familial risk of mood disorders as they develop Major Depressive Disorder. Biol. Psychiatry 78, 58–66. https://doi.org/10.1016/j.biopsych.2014.10.018

Papmeyer, M., Sussmann, J.E., Stewart, T., Giles, S., Centola, J.G., Zannias, V., Lawrie, S.M., Whalley, H.C., Mcintosh, A.M., 2016. Prospective longitudinal study of subcortical brain volumes in individuals at high familial risk of mood disorders with or without subsequent onset of depression. Psychiatry Res. Neuroimaging 248, 119–125. https://doi.org/10.1016/j.pscychresns.2015.12.009

Pedregosa, F., Varoquaux, G., Gramfort, A., Michel, V., Thirion, B., Grisel, O., Blondel, M., Prettenhofer, P., Weiss, R., Dubourg, V., Vanderplas, J., Passos, A., Cournapeau, D., Brucher, M., Perrot, M., Duchesnay, É., 2011. Scikit-learn: Machine Learning in Python. J. Mach. Learn. Res. 12, 2825−2830.

Phillips, M.L., Ladouceur, C.D., Drevets, W.C., 2008. A neural model of voluntary and automatic emotion regulation: implications for understanding the pathophysiology and neurodevelopment of bipolar disorder. Mol. Psychiatry 13, 829–857. https://doi.org/10.1038/mp.2008.65.A

Rizzo, L.B., Costa, L.G., Mansur, R.B., Swardfager, W., Belangero, S.I., Grassi-Oliveira, R., McIntyre, R.S., Bauer, M.E., Brietzke, E., 2014. The theory of bipolar disorder as an illness of accelerated aging: Implications for clinical care and research. Neurosci. Biobehav. Rev. 42, 157–169. https://doi.org/10.1016/j.neubiorev.2014.02.004

Scahill, R.I., Frost, C., Jenkins, R., Whitwell, J.L., Rossor, M.N., Fox, N.F., 2003. A Longitudinal Study of Brain Volume Changes in Normal Aging Using Serial Registered Magnetic Resonance Imaging. Arch. Neurol. 60, 989–994.

Shaw, P., Kabani, N.J., Lerch, J.P., Eckstrand, K., Lenroot, R., Gogtay, N., Greenstein, D., Clasen, L., Evans, A., Rapoport, J.L., Giedd, J.N., Wise, S.P., 2008. Neurodevelopmental trajectories of the human cerebral cortex. J. Neurosci. 28, 3586–3594. https://doi.org/10.1523/JNEUROSCI.5309-07.2008

Sibille, E., 2013. Molecular aging of the brain, neuroplasticity, and vulnerability to depression and other brain-related disorders. Dialogues Clin. Neurosci. 15, 53–65.

Smith, S.M., Vidaurre, D., Alfaro-Almagro, F., Nichols, T.E., Miller, K.L., 2019. Estimation of Brain Age Delta from Brain Imaging. Neuroimage 200, 528–539. https://doi.org/10.1101/560151

Smoller, J.W., Finn, C.T., 2003. Family, twin, and adoption studies of bipolar disease. Am. J. Med. Genet. Part C Semin. Med. Genet. 123C, 48–58. https://doi.org/10.1007/s11920-002-0046-1

Sotiras, A., Resnick, S.M., Davatzikos, C., 2015. Finding imaging patterns of structural covariance via Non-Negative Matrix Factorization. Neuroimage 108, 1–16. https://doi.org/10.1016/j.neuroimage.2014.11.045

Sotiras, A., Toledo, J.B., Gur, R.E., Gur, R.C., Satterthwaite, T.D., Davatzikos, C., 2017. Patterns of coordinated cortical remodeling during adolescence and their associations with functional specialization and evolutionary expansion. Proc. Natl. Acad. Sci. 114, 3527–3532. https://doi.org/10.1073/pnas.1620928114

Spear, L.P., 2000. The adolescent brain and age-related behavioral manifestations, Neuroscience and Biobehavioral Reviews.

Sprooten, E., Sussmann, J.E., Clugston, A., Peel, A., McKirdy, J., Moorhead, T.W.J., Anderson, S., Shand, A.J., Giles, S., Bastin, M.E., Hall, J., Johnstone, E.C., Lawrie, S.M., McIntosh, A.M., 2011. White matter integrity in individuals at high genetic risk of bipolar disorder. Biol. Psychiatry 70, 350–356. https://doi.org/10.1016/j.biopsych.2011.01.021

Tamnes, C.K., Østby, Y., Fjell, A.M., Westlye, L.T., Due-Tønnessen, P., Walhovd, K.B., 2010. Brain maturation in adolescence and young adulthood: Regional age-related changes in cortical thickness and white matter volume and microstructure. Cereb. Cortex 20, 534–548. https://doi.org/10.1093/cercor/bhp118

Tipping, M.E., 2001. Sparse Bayesian Learning and the Relevance Vector Machine. J. Mach. Learn. Res. 211–244.

Varikuti, D.P., Genon, S., Sotiras, A., Schwender, H., Hoffstaedter, F., Patil, K.R., Jockwitz, C., Caspers, S., Moebus, S., Amunts, K., Davatzikos, C., Eickhoff, S.B., 2018. Evaluation of non-negative matrix factorization of grey matter in age prediction. Neuroimage 173, 394–410. https://doi.org/10.1016/j.neuroimage.2018.03.007

Whalley, H.C., Sussmann, J.E., Romaniuk, L., Stewart, T., Kielty, S., Lawrie, S.M., Hall, J., McIntosh, A.M., 2015. Dysfunction of emotional brain systems in individuals at high risk of mood disorder with depression and predictive features prior to illness. Psychol. Med. 45, 1207–1218. https://doi.org/10.1017/S0033291714002256

Whittle, S., Lichter, R., Dennison, M., Vijayakumar, N., Schwartz, O., Byrne, M.L., Simmons, J.G., Pantelis, C., Allen, N.B., 2014. Structural brain development and depression onset during adolescence: A prospective longitudinal study. Am. J. Psychiatry 171, 564–571.

Wierenga, L.M., Langen, M., Oranje, B., Durston, S., 2014. Unique developmental trajectories of cortical thickness and surface area. Neuroimage 87, 120–126. https://doi.org/10.1016/j.neuroimage.2013.11.010

Wierenga, L.M., van den Heuvel, M.P., van Dijk, S., Rijks, Y., de Reus, M.A., Durston, S., 2016. The development of brain network architecture. Hum. Brain Mapp. 37, 717–729. https://doi.org/10.1002/hbm.23062

Wolkowitz, O.M., Reus, V.I., Mellon, S.H., 2011. Of sound mind and body: Depression, disease, and accelerated aging. Dialogues Clin. Neurosci. 13, 25–39.

World Health Organization, 2017. Depression and Other Common Mental Disorders: Global Health Estimates.

