## Supplementary Materials for "Longitudinal trajectories of brain age in young individuals at familial risk of mood disorder"

**INDEX**

Appendix A: Supplementary Methods

A.1 *Participants: Relatedness within the sample*

A.2 *Participants: Exclusions based on other or unclear diagnosis*
A.3 *Participants: Completeness of cases (Table S1)*
A.4 *MRI pre-processing: CAT toolbox and scan quality assurance*
A.5 *Brain-PAD model: Only including C-well in training sample*
A.6 *Brain-PAD model: Age distributions within samples (Figure S1)*

A.7 *Brain-PAD model: Residuals approach (Figure S2)*

Appendix B: Supplementary Results
 B.1 *Demographics: Differences in relation to attrition (Table S2)*	B.2 *Model evaluation: Discussion (Table S3)*

B.3 *Model evaluation: Brain components (Figure S3)*

B.4 *Exploratory results: The Age*Group interaction effect on Brain-PAD trajectory
 (Figure S4, Table S4)*

B.5 *Exploratory results: Brain-PAD trajectory for C-MD (Table S5, Figure S5)*

Appendix C: Supplementary References

**APPENDIX A: SUPPLEMENTARY METHODS**

**A.1 Participants: Relatedness within the sample**

The total sample (*n* = 215) included 40 participants (18.6%) who were related to at least one other participant as either siblings or cousins. More specifically, the sample included seventeen pairs and one sextet of related individuals, and two-third of these individuals (including the sextet) were with family history. One-third of the related individuals developed a mood disorder (*n* = 9 for HR-MD, *n* = 4 for C-MD). Related individuals typically experienced the same outcome; only in one case one individual of a pair developed a mood disorder, while the other remained well.

**A.2 Participants: Exclusions based on other or unclear diagnosis**
Five participants were excluded from our analysis following unclear diagnosis, or another diagnosis without mood disorder co-morbidity:

- One control participant was excluded because a follow-up SCID assessment indicated that this individual had developed Obsessive-Compulsive Disorder.
- One high-risk participant was excluded because one of the follow-up SCID assessments indicated that this individual had developed single episode psychosis without co-morbid mood disorder psychopathology.
- One high-risk participant was excluded because information from the General Practitioner indicated that the individual had experienced two episodes of hypomania (potentially related to drug use).
- Two high-risk participants were excluded because information from the General Practitioner indicated the presence of ‘low mood’, and this phrasing was considered to be too ambiguous as to whether this referred to a subclinical or clinical level of depression.

**A.3 Participants: Completeness of cases**

Table S1*.* Proportion of participants in well-groups with clinical information available per timepoint.

| **Clinical assessment** | **Years since baseline** | **C-well** | **HR-well** |
| --- | --- | --- | --- |
| Timepoint 1 | 0 | 93 (100%) | 74 (100%) |
| Timepoint 2 | 2 | 86 (92%) | 74 (100%) |
| Timepoint 3 | 4 | 53 (57%) | 54 (73%) |
| Timepoint 4 | 6 | 27 (29%) | 17 (23%) |

*Note: Participants were considered well in the absence of evidence indicating the contrary. Only imaging data from time-points 1 and 2 were used in the brain-PAD analysis. Inclusion criteria stated that all participants were well at baseline (timepoint 1), and exclusion criteria stated the exclusion of participants at familial risk (HR) for whom diagnostic information was unavailable at follow-up (timepoint 2).*
C-well, group of participants without family history who remained well; HR-well, group of participants at high familial risk who remained well.

**A.4 MRI pre-processing: CAT toolbox and scan quality assurance**
 For our study we performed cross-sectional segmentation using the Computational Anatomy Toolbox (CAT) toolbox version 12 (Gaser and Dahnke, 2018), using default settings. Scans were not segmented longitudinally in order to maximise training sample size and to keep the pre-processing pipeline consistent across those participants with and without a timepoint 2 scan. With default settings for segmentation, the CAT12 toolbox applies internal interpolation and Spatial-adaptive Non-Local Means (SANLM) (Manjón et al., 2010) as a pre-processing step in order to reduce noise normalisation, before the T1-weighted MRI scans are normalised to Montreal Neurological Institute (MNI) space using affine and non-linear registration by integration of Diffeomorphic Anatomic Registration through Exponentiated Lie algebra algorithm (DARTEL) (Ashburner, 2007) and Geodesic Shooting Normalisation (Ashburner and Friston, 2011). Segmentation is achieved by implementation of Adaptive Maximum A Posterior (AMAP) segmentation and Partial Volume Estimation (PVE) (Tohka et al., 2004). As a part of AMAP, Classical Markov Random Field (MRF) (Rajapakse et al., 1997) includes spatial information of adjacent voxels in the segmentation estimation.
 The software automatically provides a segmentation quality assurance in the form of percentage rating points (range 0-100%) for resolution, noise and bias, and a weighted average of these three components. A score above 70% is considered satisfactory. Our quality standards excluded segmentations with weighted average scores below 70% and/or resolution, noise or bias scores below 65% from analysis. In addition, we visually inspected the image quality of raw scans. We found that the CAT12 threshold was sufficient to exclude all scans with major motion-related artefacts. However, following visual inspection two additional scans were excluded because of apparent distortions in the magnetic field. In cases for which the timepoint 1 scan of the participant was missing (n = 5) or of insufficient quality (n = 7), but the timepoint 2 scan was available and the individual was known to have remained well until timepoint 2, we considered the follow-up scan as baseline scan.”

**A.5 Brain-PAD model: Only including C-well in training sample**
 A model trained on a training sample consisting only of control participants without family history of mood disorder and who remained well over the course of the study (n = 94) was devised and tested. Within this “control model” training sample, we also balanced across timepoint 1 and timepoint 2 scans in order to increase the age range of the sample. Implementation of Principal Component Analysis within the “control model” resulted in a dimensionality reduction to 76 components, as indicated by the Kaiser Criterion (eigenvalue > 1). Subsequently, Relevance Vector Regression with a linear kernel was applied, similarly as within the eventually implemented model. For a fair comparison, the “control model” was evaluated based on the brain age predictions for the eventually used training sample: that is 167 observations from C-well and HR-well, balanced across timepoint 1 and timepoint 2 measurements. In this way, the evaluation of the “control model” corresponded to the evaluation of the eventually implemented model.
 Results of the “control model” evaluation showed a significant but small Pearson correlation between brain age predictions and corresponding chronological ages (*r*(165) = .25*, p* = .001) and a Mean Absolute Error (MAE) of 2.32 years. The average brain age prediction was 0.20 years lower than the average chronological age measure, showing a slight underestimation in brain age prediction. The relatively low correlation between brain age prediction and chronological age in combination with the small underestimation in the model reflect its suboptimality, leading to the additional inclusion of HR-well participants (*n* = 74) in the training sample of our eventually implemented model. Following this change in training sample, evaluation of the eventually implemented model showed an improvement in accuracy reflected by a medium correlation (Pearson correlation, *r*(165) = .40, *p* < .001) between brain age predictions and chronological age, a slightly lower MAE of 2.21 years, and no under- or overestimation (0.00 years) in brain age predictions – hence its implementation.
 As a consequence of training on all individuals who remained well, brain age predictions were slightly biased towards the neuroanatomy of HR-participants. As HR-participants are potentially showing slight deviations from normative brain development, normative development is most accurately reflected by brain-PAD values of the control group (instead of brain-PAD = 0).

**A.6 Brain-PAD model: Age distributions within samples**

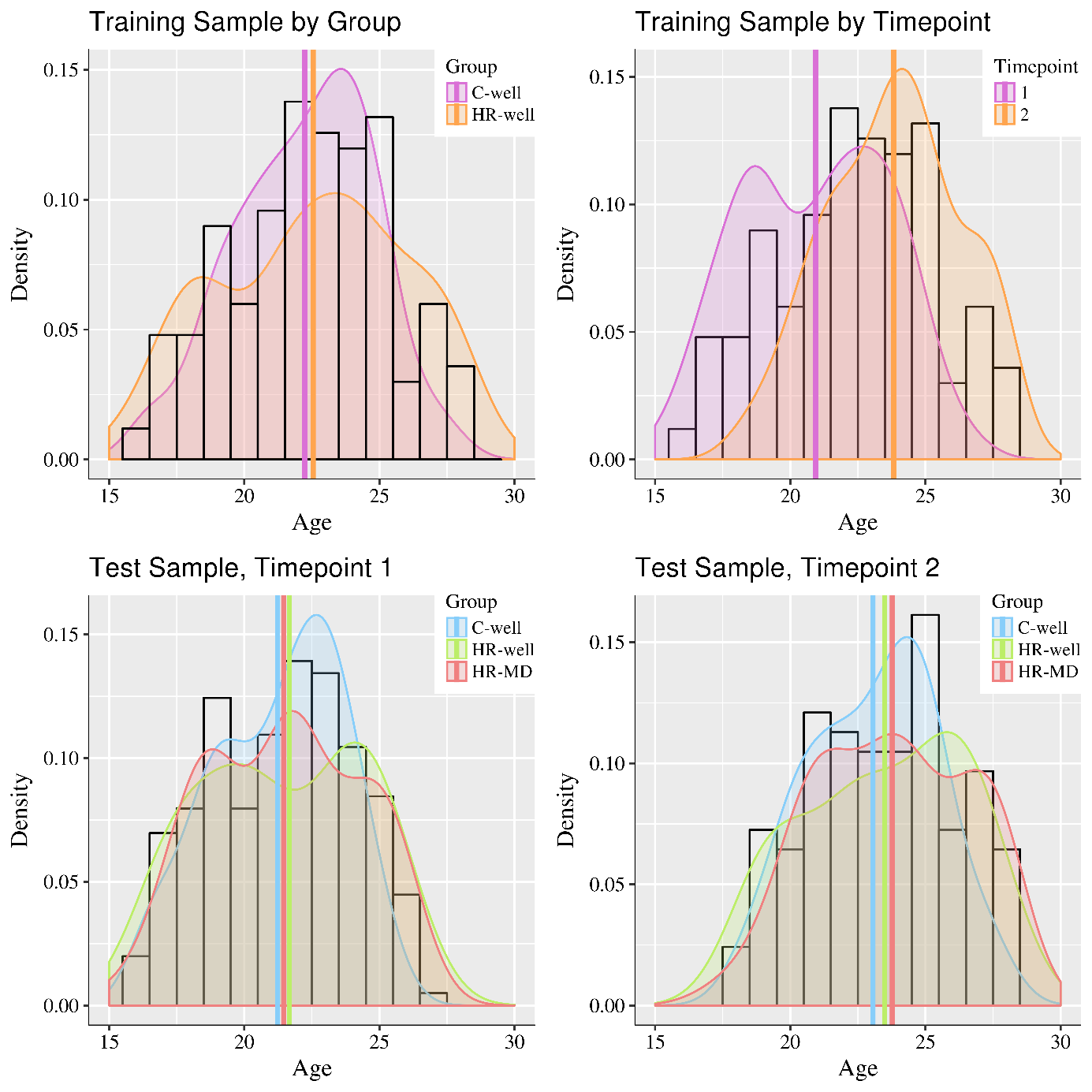

*Figure S1.* Density plots visualising age distributions of the samples. In all figures, white histograms represent the total sample, whereas the coloured line graphs and shaded areas display density plots of subsamples. Coloured vertical lines display mean ages per subsample. In the upper row, density plots represent the age distribution of the training sample per group (upper left) and per timepoint (upper right). In the lower row, density plots represent the age distributions of the test samples per group for timepoint 1 (lower left) and timepoint 2 (lower right).

C-well, group of participants without family history who remained well; HR-MD, group of participants at high familial risk who developed a mood disorder; HR-well, group of participants at high familial risk who remained well.

**A.7 Brain-PAD model: Residuals approach**

A residuals approach was applied by calculating the brain-predicted age difference (brain-PAD) based on the residuals from the regression model that regresses brain age prediction on chronological age and sex within the training sample. In other words, we fit a linear regression model (‘brain age prediction ~ chronological age + gender’) on the training sample data which resulted in two regression lines corresponding to each gender (Figure S2). Subsequently, for each participant the expected brain age prediction was calculated as the point on the regression line corresponding to the chronological age and gender of the participant. Brain-PAD was calculated by subtracting this expected prediction from the actual brain age as predicted by the brain age prediction model. This corresponds to the distance (‘residual’) of the participant’s observation to the training sample regression line within a scatter plot of chronological age against brain age prediction (see Figure S1). The brain-PAD residuals approach takes into account the inaccuracies of model predictions by using the observed relationship between brain age prediction and chronological age, therefore sufficiently addressing the underfitting of our model (Smith et al., 2019). Consequently, brain-PAD calculations are unbiased and uncorrelated to chronological age, *r*(166) = 0.00, *p* = 1) and as such more reliable than a simple subtraction method.

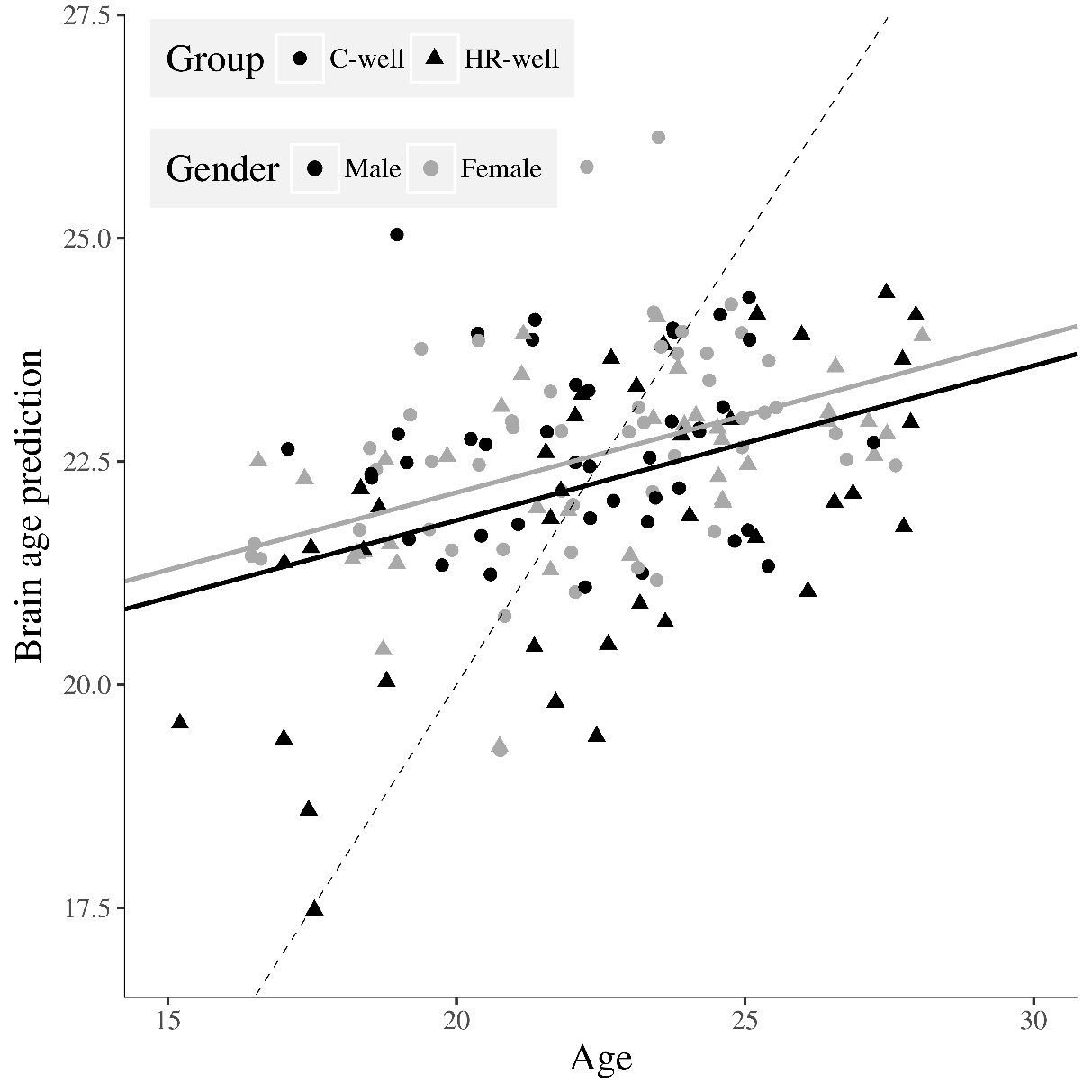

*Figure S2.* Scatterplot for the training sample that shows brain age prediction against chronological age. The solid black line graph (brain age prediction = 18.38 + 0.17 * chronological age) and grey line graph (brain age prediction = 18.69 + 0.17 * chronological age) represent the regression lines for males and females respectively. These regression formulas were used in the residuals approach to determine the expected brain age prediction for each participant. The result was subsequently subtracted from the actual brain age prediction to calculate the brain-predicted age difference (brain-PAD). The dotted line is a diagonal reference line (for brain age prediction = chronological age).

**APPENDIX B: SUPPLEMENTARY RESULTS**

**B.1 Demographics: Differences in relation to attrition**

Table S2. Participants demographic and clinical information in relation to attrition to follow-up.

|  | **Follow-up**  *n =* 124 | **No follow-up**  *n* = 78 | ***p*-value** |
| --- | --- | --- | --- |
| Age, timepoint 1^a^ | 21.4 (4.4) | 21.7 (4.3) | .79 |
| Gender^b^ |  |  | .95 |
| *Male* | 43 (45.7%) | 37 (50.6%) |  |
| *Female* | 51 (54.3%) | 36 (49.4%) |  |
| Handedness^b^ |  |  | .34 |
| *Left* | 9 (7.3%) | 5 (6.4%) |  |
| *Right* | 114 (91.9%) | 70 (89.7%) |  |
| *Mixed* | 1 (0.8%) | 1 (1.3%) |  |
| *Unknown* | 0 (0.0%) | 2 (2.6%) |  |
| NART score^a^ | 111 (8.8) | 110 (11.0) | .59 |
| HRSD, timepoint 1^a^ | 0 (2.0) | 0 (2.0) | .85 |

Note: Individual NART scores were averaged over all completed assessments (max. 4).

^a^ Medians and interquartile ranges for variables not normally distributed (Kruskal-Wallis test)

^b^ Frequency and percentages for categorical variables (chi-squared test).

HRSD, Hamilton Rating Scale for Depression; NART, National Adult Reading Test.

**B.2 Model evaluation: Discussion**

The model implemented within the current study shows a medium correlation (Pearson correlation, *r*(165) = .40, *p* < .001) between brain age prediction and chronological age within the training sample, and a mean absolute error (MAE) of 2.21 years. Dividing the MAE by the training sample age range of 15.2-28.1 years, the scaled MAE corresponds to 0.17 years. Table S3 provides the MAE and correlation measures per included subsample for more thorough evaluation of model performance.

Table S3. Model evaluation per sample and timepoint.

| **Sample** | **Timepoint** | ***N*** | **Age range** | **MAE** | **Scaled MAE** | ***r*** |
| --- | --- | --- | --- | --- | --- | --- |
| Training  .  . | Full sample | 167 | 15.2-28.1 | 2.21 | 0.17 | 0.40*** |
|  | Timepoint 1 | 84 | 15.2-26.6 | 2.13 | 0.19 | 0.41*** |
|  | Timepoint 2 | 83 | 18.3-28.1 | 2.29 | 0.23 | 0.10 |
| Outwith training^φ^ | Full sample | 93 | 16.3-26.0 | 2.05 | 0.21 | 0.17 |
| C-well  . | Timepoint 1 | 93 | 16.3-25.6 | 2.01 | 0.22 | 0.22* |
|  | Timepoint 2 | 46 | 18.3-27.6 | 2.11 | 0.23 | 0.04 |
| HR-well  . | Timepoint 1 | 74 | 15.2-26.6 | 2.22 | 0.20 | 0.42*** |
|  | Timepoint 2 | 47 | 17.6-28.1 | 2.36 | 0.22 | 0.36* |
| HR-MD  . | Timepoint 1 | 35 | 16.0-30.0^$^ | 2.74 | 0.20^$^ | 0.20 |
|  | Timepoint 2 | 31 | 18.1-28.1 | 2.52 | 0.25 | 0.39* |

* *p < .05, ** p < .01, *** p < .001
^φ^Well-group participants (i.e. training sample) scans that were not included in the training sample.
^$^One participant was aged 30.0 at timepoint 1, because baseline scan was missing and thus follow-up scan was treated as baseline scan. Without this individual, age range is 16.0-26.1, and scaled MAE = 0.27.*

C-well, group of participants without family history who remained well; HR-MD, group of participants at high familial risk who developed a mood disorder; HR-well, group of participants at high familial risk who remained well; MAE, mean absolute error.

Previous studies with a much broader age range within their sample – including young and older participants (e.g. 18-65 years old) – generally achieve a Pearson correlation of around 0.9 and an MAE of over 4 years when predicting brain age, corresponding to a scaled MAE of ~0.10 (Cole et al., 2017; Schnack et al., 2016). Franke et al. (2012) included children and adolescents aged 4 to 18 years and achieved Pearson correlations of >0.9 in combination with MAEs ranging between 1.1 and 1.3 years within their samples.
 Although we acknowledge the relatively low correlation of our model, it is important to note that measures of model accuracy actually also reflect variability in true brain age, with greater variability producing a lower correlation between chronological age and predicted brain age, as well as higher MAE. At the juncture between the end of childhood development and the gradual onset of degeneration throughout adulthood, variability in true brain age is likely to be higher, which may partly explain our relatively low correlation and higher MAE. Furthermore, we suggest that this relatively low correlation is likely to be affected by the narrow age range within the cohort, as well as our relatively limited cohort sample size. We also note that the MAE is comparable to other studies in the field (only slightly higher, as expected given the consideration above). Therefore, we argue that the overall performance of our brain age prediction is sufficient for the aim of the current study.

**B.3 Model evaluation: Brain components**

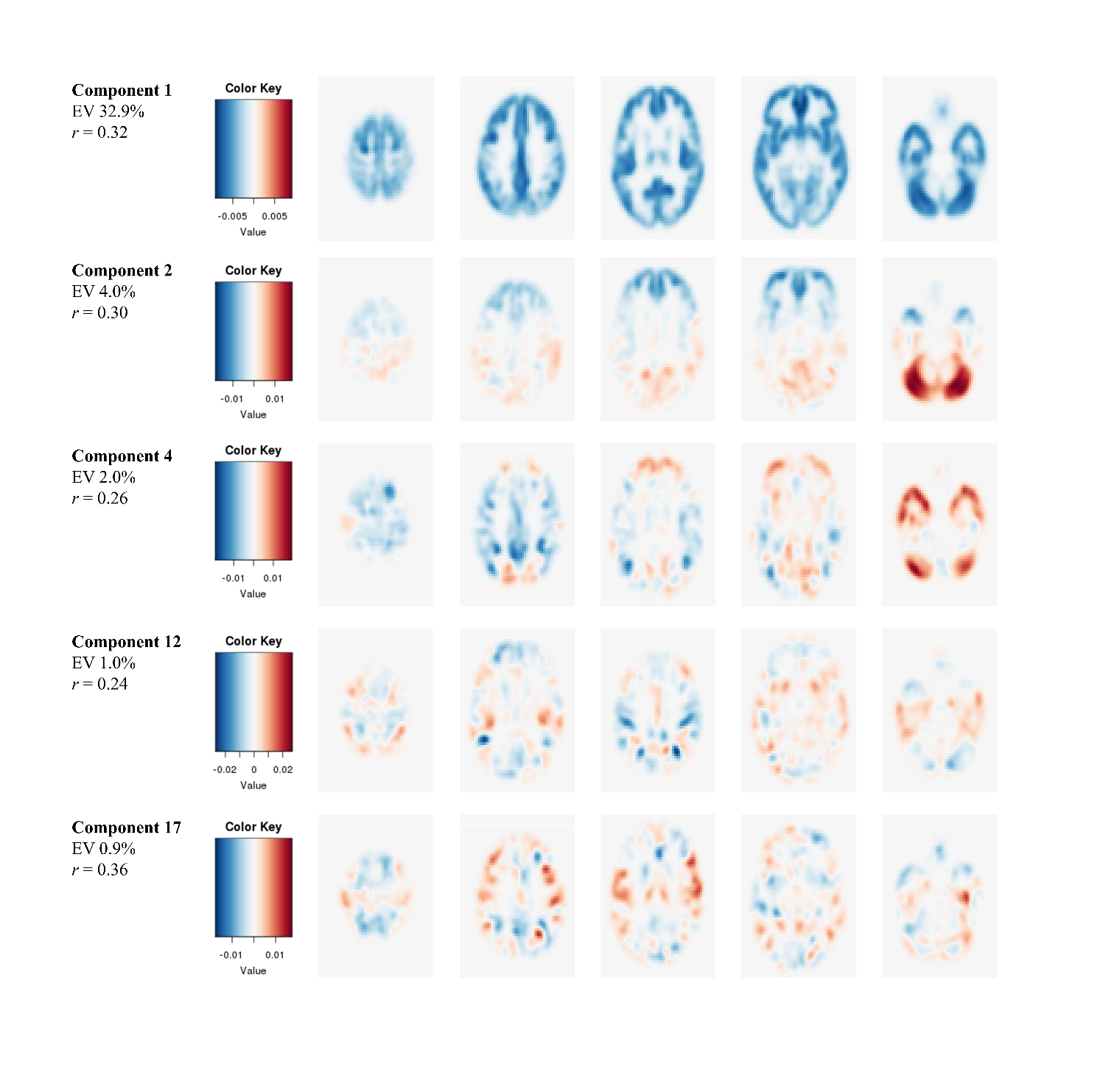

*Figure S3.* This figure graphically displays the loadings of the five PCA components that (after FDR-correction) significantly correlated with the brain age prediction in the training sample, and provides per component the statistic of this Pearson correlation (*r*) and the explained variance (EV) with respect to all inter-subject variance in grey matter. For all components, blue coloured areas reflect that lower brain grey matter volumes were associated with higher brain age predictions (negative correlation), while red coloured areas reflect associations in the opposite directions (positive correlation).
 Specifically, brain component loadings and values were retrieved for each participant within the training sample (i.e. within each iteration of the leave-one-out cross-validation loop), after which component values were correlated to the brain age prediction. For significantly correlated components we averaged over the component loadings that were extracted from each iteration (i.e. for each test sample) under the assumption that the brain components would sufficiently map onto each other.

**B.4 Exploratory results: The Age*Group interaction effect on Brain-PAD trajectory**

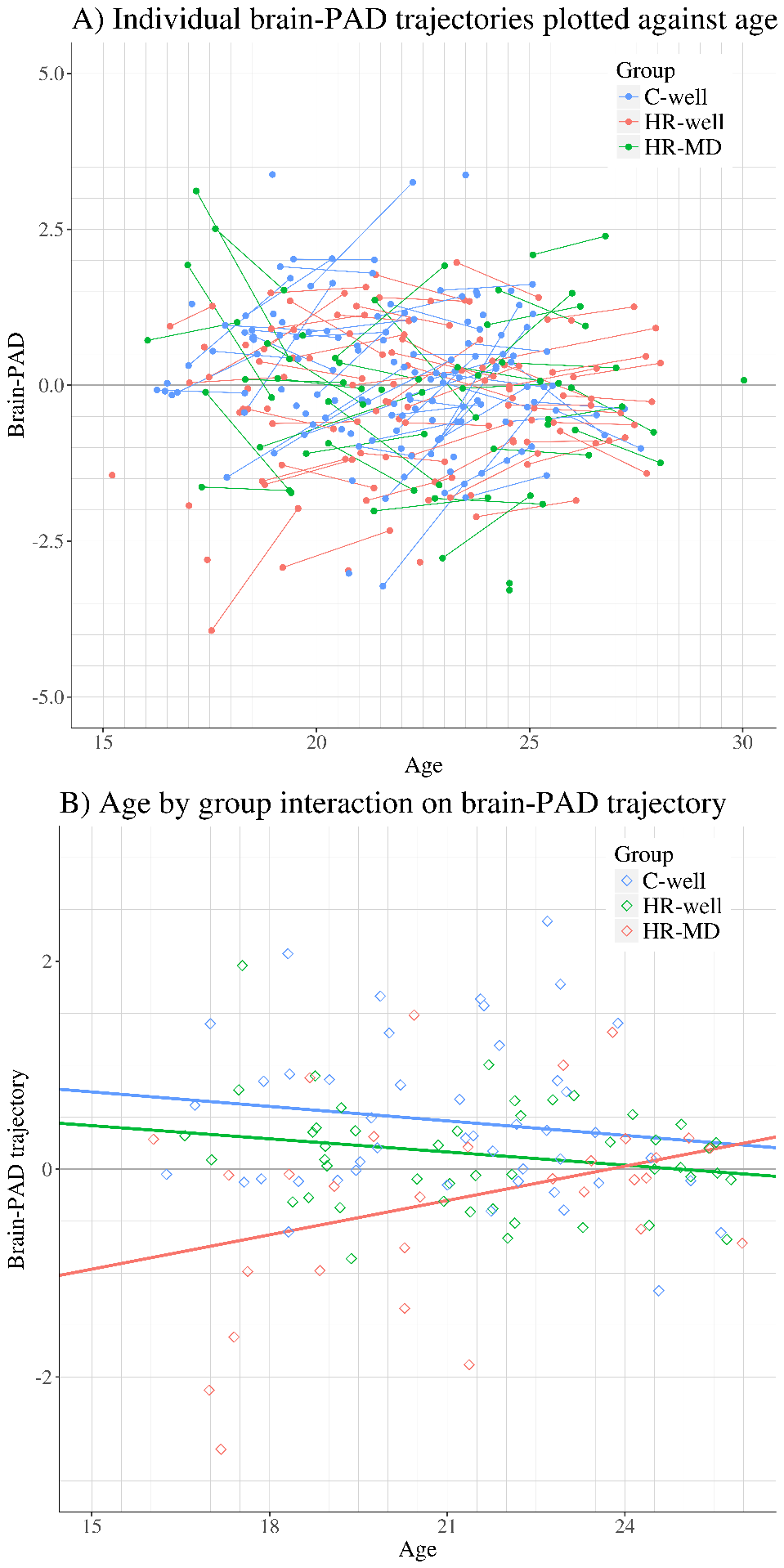

*Figure S4*. Brain-PAD trajectories as a function of age.

Figure S4A (upper plot) shows brain-PAD trajectories of each individual, coloured by group; each dot represents an assessment at either timepoint, and two measurements of one individual are connected with a line graph in order to visualise the trajectory.

Figure S4B (lower plot) shows the age by group interaction effect on brain-PAD trajectory. Each dot represents the trajectory of one individual, i.e. the difference in brain-PAD between the two assessments, and is coloured by group. The coloured line graphs show the average brain-PAD trajectory per group as a function of age at baseline.

Table S4. Fixed effects of the longitudinal linear mixed model investigating the age by group interaction on brain-predicted age difference (brain-PAD) trajectory.

| **Longitudinal model** | | | | | |
| --- | --- | --- | --- | --- | --- |
| **Fixed effect** | ***β-*coefficient** | ***SE*** | ***df*** | **t-value** | ***p*-value** |
| (Intercept) | 0.38 | 0.14 | 107 | 2.74 | .007** |
| Age (at baseline) | -0.16 | 0.16 | 11 | -1.02 | .33 |
| HR-well | -0.38 | 0.20 | 11 | -1.92 | .06 |
| HR-MD | -0.90 | 0.22 | 11 | -4.13 | .002** |
| Age*HR-well | 0.01 | 0.20 | 11 | 0.07 | .94 |
| Age*HR-MD | 0.53 | 0.22 | 11 | 2.48 | .03* |

** p < .05, ** p < .01, *** < .001*
 *Note: results were referenced by the control group participants who remained well (C-well). β-coefficients are standardised following scaling of the outcome variable ‘brain-predicted age difference (brain-PAD) trajectory’ and predictor ‘age at baseline’. Inclusion of ‘chronological ageing’ (difference in age between baseline and follow-up) as covariate yielded similar results.*

C-well, group of participants without family history who remained well; HR-MD, group of participants at high familial risk who developed a mood disorder; HR-well, group of participants at high familial risk who remained well.

**Summary of exploratory findings**

Figure S4B suggest an age by group interaction effect for HR-MD, in which younger individuals who subsequently develop a mood disorder show a greater deceleration in brain-PAD between baseline and follow-up. The results of our exploratory longitudinal model (Table S4) indeed show a statistically significant age by group interaction effect on brain-PAD trajectory for the HR-MD group, which is not present in the HR-well group. Furthermore, brain-PAD trajectories appear to stabilise within older individuals. In other words, on average the brain-predicted age difference (brain-PAD) barely changed between the two timepoints within individuals >21 years at baseline. One implication of these findings is that for future research it may be most useful to investigate brain-PAD trajectories in younger individuals (e.g., 14-21 years old).

**B.5 Exploratory results: Brain-PAD trajectory for C-MD**

Table S5. Fixed effects of the linear mixed model applied to investigate brain-predicted age difference (brain-PAD) within the group of participants without family history who developed a mood disorder (C-MD).

| **Fixed effect** | ***β-*coefficient** | ***SE*** | ***df*** | **t-value** | ***p*-value** |
| --- | --- | --- | --- | --- | --- |
| C-MD | 0.35 | 0.36 | 25 | 0.98 | .34 |
| Timepoint 2*C-MD | -0.44 | 0.26 | 129 | -1.67 | .10 |

*Note: Only results regarding C-MD are presented within the table. Results were referenced by the control group participants who remained well (C-well). β-coefficients are standardised following scaling of the outcome variable brain-predicted age difference (brain-PAD).*

C-MD, group of participants without family history who developed a mood disorder.

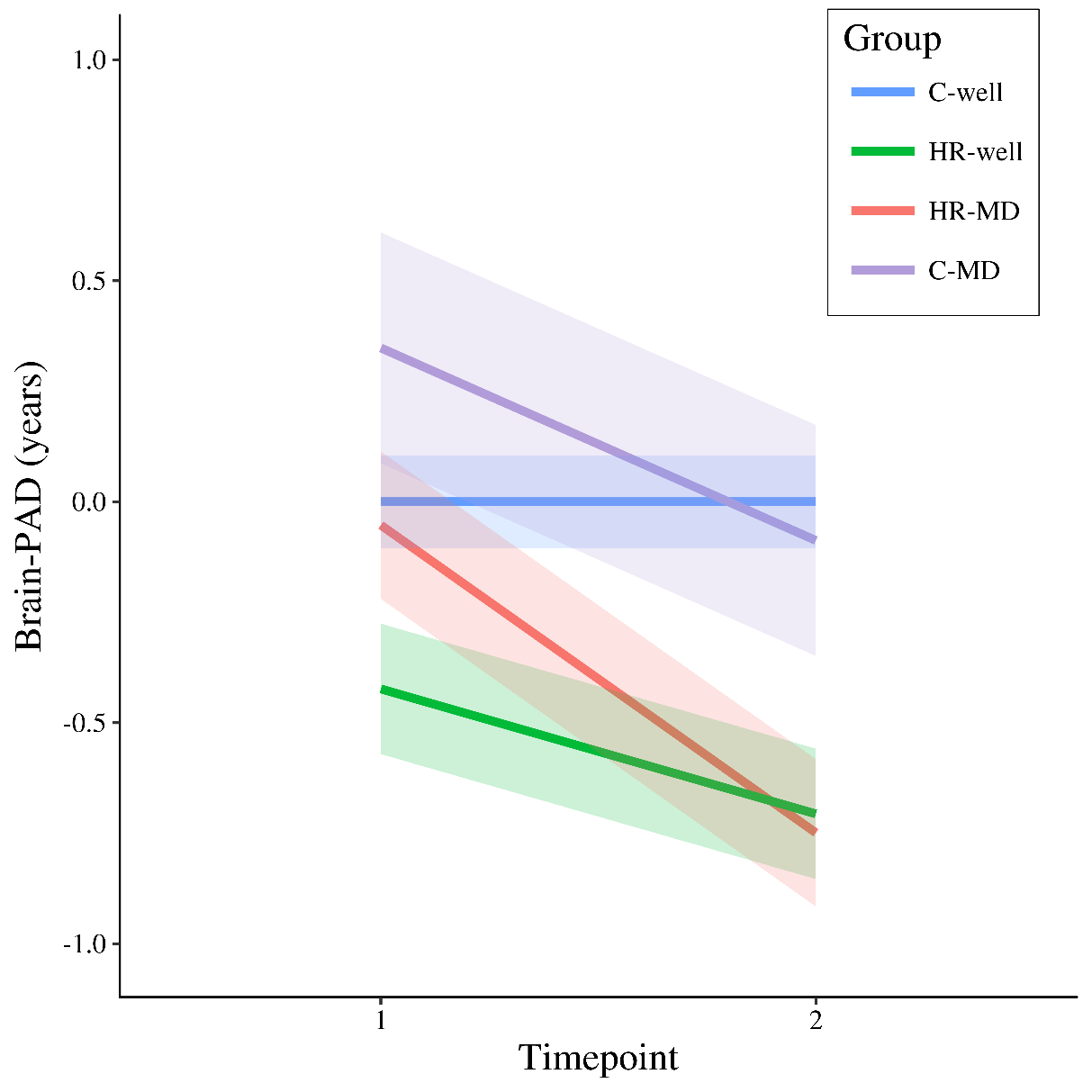

*Figure S5.* Modelled fixed effects of the brain-predicted age difference (brain-PAD) per group, including C-MD, for clarity corrected for effects in C-well (i.e., the intercept and timepoint coefficient) as this group functions as control and reference group within the main analysis. Shaded areas display standard errors of the timepoint fixed effects per group.

Brain-PAD, brain-predicted age difference; C-MD, group of participants without family history who developed a mood disorder; C-well, group of participants without family history who remained well; HR-MD, group of participants at high familial risk who developed a mood disorder; HR-well, group of participants at high familial risk who remained well.

**Appendix C: Supplementary References**

Ashburner, J., 2007. A fast diffeomorphic image registration algorithm. Neuroimage 38, 95–113. https://doi.org/10.1016/j.neuroimage.2007.07.007

Ashburner, J., Friston, K.J., 2011. Diffeomorphic registration using geodesic shooting and Gauss – Newton optimisation. Neuroimage 55, 954–967. https://doi.org/10.1016/j.neuroimage.2010.12.049

Cole, J.H., Poudel, R.P.K., Tsagkrasoulis, D., Caan, M.W.A., Steves, C., Spector, T.D., Montana, G., 2017. Predicting brain age with deep learning from raw imaging data results in a reliable and heritable biomarker. Neuroimage 163, 115–124. https://doi.org/10.1016/j.neuroimage.2017.07.059

Franke, K., Luders, E., May, A., Wilke, M., Gaser, C., 2012. Brain maturation: Predicting individual BrainAGE in children and adolescents using structural MRI. Neuroimage 63, 1305–1312. https://doi.org/10.1016/j.neuroimage.2012.08.001

Gaser, C., Dahnke, R., 2018. CAT – A Computational Anatomy Toolbox for SPM.

Manjón, J. V, Coupé, P., Martí-Bonmatí, L., Collins, D.L., Robles, M., 2010. Adaptive Non-Local Means Denoising of MR Images With Spatially Varying Noise Levels. J. Magn. Reson. Imaging 31, 192–203. https://doi.org/10.1002/jmri.22003

Rajapakse, J.C., Giedd, J.N., Rapoport, J.L., 1997. Statistical approach to segmentation of single-channel cerebral MR images. IEEE Trans. Med. Imaging 16, 176–186.

Schnack, H.G., van Haren, N.E.M., Nieuwenhuis, M., Hulshoff Pol, H.E., Cahn, W., Kahn, R.S., 2016. Accelerated brain aging in Schizophrenia: A longitudinal pattern recognition study. Am. J. Psychiatry 173, 607–616. https://doi.org/10.1176/appi.ajp.2015.15070922

Smith, S.M., Vidaurre, D., Alfaro-Almagro, F., Nichols, T.E., Miller, K.L., 2019. Estimation of Brain Age Delta from Brain Imaging. Neuroimage 200, 528–539. https://doi.org/https://doi.org/10.1101/560151

Tohka, J., Zijdenbos, A., Evans, A., 2004. Fast and robust parameter estimation for statistical partial volume models in brain MRI. Neuroimage 23, 84–97. https://doi.org/10.1016/j.neuroimage.2004.05.007
